# Medulloblastoma-associated DDX3X mutants are oncogenic having a defect in translation-promoting activity but functional in stress granule formation and interferon signaling

**DOI:** 10.1101/2023.08.15.553361

**Authors:** Satishkumar Vishram Singh, Harish Shrikrishna Bharambe, Shalaka Arun Masurkar, Purna Bapat, Nikhil Gadewal, Neelam Vishwanath Shirsat

## Abstract

DDX3X, a DEAD box-containing RNA helicase, is known to play diverse roles in RNA metabolism, stress response, innate immunity, and cancer. Medulloblastoma is the single most common malignant brain tumor in children. DDX3X is recurrently mutated in the WNT and SHH subgroups of medulloblastoma. CRISPR-Cas9 mediated DDX3X knockout was successful in the HEK293FT cells but generated only non-truncating indels in the medulloblastoma cells suggesting DDX3X is necessary for the viability of the cells. Downregulation of DDX3X expression using shRNA also brought about a considerable reduction in proliferation, clonogenic potential, and anchorage-independent growth of the medulloblastoma cells. Thus, DDX3X expression was found to be essential for the survival, growth, and malignant potential of the medulloblastoma cells consistent with the non-truncating nature of medulloblastoma-associated DDX3X mutations. The medulloblastoma-associated DDX3X mutants were found to be defective in their ability to drive the translation of mRNAs with complex 5’-UTR that is dependent on the ATP-dependent helicase activity of DDX3X. These helicase defective DDX3X mutants could restore the expression of interferon signaling genes and malignant potential lost upon DDX3X knockdown in medulloblastoma cells. Their N-terminal domain is intact and was found to be functional in stress granule formation. DDX3X mutants upregulated expression of malignancy-related genes suggesting tumor suppressive role for the helicase activity in the medulloblastoma pathogenesis. Inhibitors of the N-terminal domain of DDX3X which is essential for the viability of medulloblastoma cells could have therapeutic potential in the treatment of WNT and SHH subgroup medulloblastomas.

## Introduction

DDX3X is an RNA helicase that belongs to a highly conserved subfamily of the DEAD (Asp-Glu-Ala-Asp) box containing RNA helicases (1). DDX3X is abundantly expressed in all tissues and plays diverse roles in RNA metabolism, innate immune response, cell cycle, and cellular stress response (2,3). DDX3X plays a role in mRNA transcription, splicing, translation, and transport. It is shown to be a critical component of the TBK1/IKKε mediated induction of the anti-viral interferon I signaling pathway and is also involved in the inflammasome formation (4,5).

Mutations in DDX3X have been reported in several cancers (6). DDX3X is the most frequently mutated gene in Natural Killer/ T cell lymphoma (7). Recurrent truncating mutations in DDX3X are also reported in chronic lymphocytic leukemia suggesting a tumor suppressor role for DDX3X (8). On the other hand, DDX3X is reported to play an oncogenic role in breast cancer, colorectal cancer, and lung cancer cells by promoting epithelial-mesenchymal transition and WNT signaling (9). Thus, DDX3X appears to play a dual role in the pathogenesis of cancer.

Medulloblastoma is the single most common brain tumor in children (10). It is a highly malignant tumor that consists of four molecular subgroups called WNT, SHH, Group 3, and Group 4. WNT and SHH subgroup medulloblastomas are primarily driven by the canonical WNT and Sonic hedgehog signaling pathway, respectively. Interestingly, mutations in DDX3X are restricted to only these two subgroups (11,12). DDX3X is the second most mutated gene after the *CTNNB1* gene encoding beta-catenin in the WNT subgroup medulloblastomas. DDX3X mutations occur in ∼10% of pediatric and 50% of adult SHH subgroup medulloblastomas. Almost all DDX3X mutations in medulloblastoma are non-truncating missense mutations occurring in the two RecA domains involved in the helicase activity of the protein. Only two truncating DDX3X mutations have been reported so far, at the amino acid position 242 and 426 of the 662 amino acid long protein (13,14). The truncating mutation at the amino acid 242 position lacks the RecA domains required for the helicase activity but keeps its N-terminal domain intact (13). Some of the missense mutations have been reported to be defective in RNA-stimulated ATP hydrolysis suggesting loss of ATP-dependent helicase activity (15) and some mutants which could be tested in vitro, were found to be defective in the helicase activity (16).

The present study was carried out to understand the role of DDX3X and its medulloblastoma-associated mutants in the pathogenesis of medulloblastoma and identify the underlying molecular mechanism.

## Materials and Methods

### Cell culture

Medulloblastoma cell line Daoy was procured from the American Type Culture Collection, Manassas, VA, USA. Human embryonic kidney cell line HEK293FT was obtained from Thermo Fisher Scientific, Waltham, MA, USA. The authenticity of the cell lines was validated by the short tandem repeat marker profiling and the cells were periodically checked for mycoplasma contamination (17). The cell lines were grown in Dulbecco’s Modified Eagle Medium containing 10% fetal calf serum at 37°C in a moisturized CO2 incubator.

### Generation of DDX3X knockout clones

CRISPR-Cas9 technology was used to generate DDX3X knockout clones of HEK293FT and Daoy cell lines. Guide RNAs targeting the DDX3X gene were designed in exon 1 and exon 4 of the DDX3X gene using the gRNA designing tools (http://crispr.mit.edu/) (Supplementary Table 1). The oligonucleotides corresponding to the gRNA sequences were cloned in the pLentiCRISPRv1 vector (Addgene Plasmid # 49535), a doxycycline-inducible lentiviral vector. The cells were transduced with the lentiviral particles carrying the gRNA and clones were selected in the presence of puromycin. The clones were screened for indels using a PCR-based strategy (18), wherein two pairs of primers were designed with one primer overlapping the potential indel site next to the Protospacer Adjacent Motif (PAM) (Supplementary Table 1). The indel introduced by Cas9 would affect only one of the two amplicons. The clones identified as having indels in PCR analysis were then checked for DDX3X knockout by the western blotting and validated by Sanger sequencing.

### ShRNA mediated downregulation of DDX3X expression in medulloblastoma cells

DDX3X expression was downregulated using two independent shRNA oligonucleotides, one targeting the open reading frame (shORF) and the other targeting the 3’-untranslated (shUTR) region (Supplementary Table 1), cloned in a doxycycline-inducible lentiviral vector Tet-pLKO-puro vector (Addgene #21915). Daoy medulloblastoma cells were transduced with the lentiviral particles carrying the shRNA construct and stably transduced cells were selected in the presence of puromycin. DDX3X expression was checked by western blotting using an anti-DDX3X antibody (D19B4, Cell Signaling Technology, Danvers, MA, USA).

### Effect of DDX3X knockdown on the growth of medulloblastoma cells

The effect of shRNA-mediated DDX3X knockdown on the proliferation, clonogenicity, and anchorage-independent growth of the medulloblastoma cells was studied by the MTT assay, clonogenic assay, and soft agar colony forming assay, respectively. For the MTT assay,1000 cells were seeded per well of a 96-well microtiter plate with and without doxycycline treatment for 48 h. The medium was replenished at 2-day intervals. The cells were incubated with MTT for 4 h, formazan crystals formed were dissolved in acidified SDS, and the optical density (O.D) was read on an Epoch2 Microplate Reader, Bio Tek Instruments, Winooski, VT, USA. For the clonogenic assay, 1000 cells were seeded in a 55 mm tissue culture plate with and without doxycycline induction, and the colonies formed were stained using crystal violet and counted on a stereo microscope. For soft agar assay, 7500 cells were seeded in 0.3% agar on top of a 1% agar layer in a 35 mm tissue culture plate before and after 48 h doxycycline treatment. After about 7 days of incubation, colonies consisting of at least 15 -20 cells were counted on an inverted phase contrast microscope.

### Effect of mutations in DDX3X on its translation regulation activity

The effect of mutations in DDX3X on its translation regulation activity was studied by the luciferase reporter assay. The 5’UTR region of the cyclin E gene was amplified from normal human peripheral blood lymphocytes genomic DNA using the Phusion High Fidelity DNA polymerase and was cloned downstream of the SV40 promoter and upstream of the firefly luciferase cDNA sequence in the pGL3 control vector (Promega, Madison, WI, USA). The DDX3X knockout HEK293FT cells were co-transfected with the Cyclin E-5’UTR luciferase reporter construct, Enhanced Green Fluorescent Protein (EGFP) expression vector, and wild-type or mutant DDX3X expression constructs using the calcium phosphate-*N*-bis(2-hydroxyethyl)-2-amino-ethane sulfonic acid (BES) buffer method (19). The total protein was extracted from the cells 72 h after transfection. The luciferase activity of the cell extracts was measured using D-Luciferin as a substrate. The luminescence and EGFP fluorescence were measured using the Cytation 5 Hybrid Multi-mode reader (Bio Tek Instruments, Winooski, VT, USA).

### Transfection of wild-type and mutant DDX3X in DDX3X knockdown medulloblastoma cells

Human DDX3X cDNA was a kind gift from Dr. Laura Madrigal-Estebas, Trinity College, Dublin, Ireland. Full-length cDNA was cloned in p3XFLAG-CMV™-10 vector (Sigma-Aldrich, St Louis, MO, USA) such that the 3X-FLAG tag at its 5’end was in the same reading frame as DDX3X. Mutations in the helicase encoding domain of the DDX3X cDNA were introduced by the site-directed mutagenesis using the Phusion High Fidelity DNA Polymerase (New England Biolabs, Ipswich, MA, USA) by overlap extension PCR method as described (20).

Both wild-type and mutant DDX3X cDNA plasmids were transfected in the shRNA (shUTR) knockdown polyclonal population of Daoy cells using the X-tremeGENE™ HP DNA transfection reagent (Sigma-Aldrich, MO, USA). Expression of the exogenous DDX3X was checked by western blotting using an anti-FLAG® M2 antibody, (Sigma-Aldrich, MO, USA) and an anti-DDX3X antibody (D19B4, Cell Signaling Technology, Danvers, MA, USA).

### Effect of mutations in DDX3X on its role in stress granule formation

The effect of mutations in DDX3X on its role in stress granule formation was studied by immunofluorescence imaging. The stress granule formation was studied in the DDX3X vector control cells, shUTR knockdown cells, and the knockdown cells stably transfected with wild-type or mutant DDX3X of the Daoy cell line. The shRNA expression was induced with doxycycline treatment of the cells for 48 h. For induction of the stress granule formation, the cells were treated with 200 µM Arsenic trioxide for 30 min. The cells were fixed using freshly prepared 4% (w/v) formaldehyde and permeabilized using 0.3% (w/v) Triton X-100. The fixed and permeabilized cells were incubated with primary antibodies overnight at 4^°^C after blocking with 3% bovine serum albumin followed by incubation with fluorescent dye-conjugated secondary antibodies and imaged on a Leica confocal microscope. The antibodies used are listed below.

Anti-eIF4G (sc-11373, Santa Cruz Biotechnology, TX, USA); anti-FLAG® M2 antibody (Sigma-Aldrich, MO, USA); Alexa Fluor® 647 AffiniPure Donkey anti-Rabbit IgG (H+L) (711-605-152, Jackson ImmunoResearch, PA, USA) and Donkey anti-Mouse IgG H&L (Alexa Fluor® 488) (ab150105, Abcam, UK).

### Transcriptome analysis of DDX3X knockdown medulloblastoma cells before or after expression of exogenous wild-type or mutant DDX3X

Total RNA was extracted from Daoy cells, and their shRNA-mediated DDX3X knockdown polyclonal populations using the Qiagen RNeasy kit (Hilden, Germany) after 48 h doxycycline treatment. Total RNA was also extracted from the Daoy DDX3X knockdown cells transfected with the wild-type or mutant DDX3X. For transcriptome sequencing, libraries were prepared using the Illumina Hiseq TrueSeq RNA library preparation kit v2 from Illumina (San Diego, CA, USA) and were sequenced on a Hiseq X platform (Illumina, San Diego, CA, USA) to get a minimum of 10 million single-end reads per sample. The sequencing data were aligned to the reference human genome GRCh37-hg19 using the HISAT2 algorithm using default parameters (https://ccb.jhu.edu/software/hisat2). HTseq algorithm (www.bioinformatics.babraham.ac.uk) was used to get raw counts of the number of reads per transcript. The transcripts significantly differentially expressed upon the DDX3X knockdown were identified using the DESeq2 software (https://bioconductor.org/packages/release/bioc/html/DESeq2.html) in the R Bioconductor environment following the variance stabilizing normalization. The transcriptome data are deposited in the Gene Expression Omnibus database (https://www.ncbi.nlm.nih.gov/geo/query/acc.cgi?acc=GSE201967). ClueGO plug-in v2.5.9 of the Cytoscape analysis software v3.9.1 (https://cytoscape.org) was used to identify the biological pathways enriched in the genes significantly differentially expressed upon DDX3X knockdown and their interaction network was built using the GO_Biological Processes database [14]. Heat maps of the genes were plotted using the Multiple Experiment Viewer (http://mev.tm4.org) after the median centering of the data.

All experiments were performed at least three times and the statistical significance was deduced by the Student’s t-test.

## Results

To decipher the role of the *DDX3X* gene in medulloblastoma pathogenesis, the effect of mutations in *DDX3X* on the growth and malignant potential of medulloblastoma cells was investigated. DDX3X is abundantly expressed in most cell types including the medulloblastoma cell line Daoy belonging to the SHH subgroup. Therefore, the expression of endogenous DDX3X was downregulated before assessing the activity of exogenous mutant DDX3X.

### CRISPR-Cas9 mediated DDX3X knockout in medulloblastoma cells generated only non-frameshift indels

For DDX3X knockout, guide RNAs (gRNAs) were designed in exon 1 and exon 2 of the *DDX3X* gene and cloned in the pLentiCRISPRv1 vector (Addgene Plasmid # 49535). HEK293FT and Daoy cells were transduced with the pLentiCRISPR constructs carrying the gRNA and monoclonal populations were selected in the presence of puromycin. The clones were screened by PCR for the presence of indels around the targeted PAM site in the DDX3X gene (18). Several clones were identified as DDX3X knockout by PCR screening as depicted in Fig. 1A. Absence of 206 bp amplicon indicates the presence of Indel at the PAM site. Western blot analysis of HEK293FT clones showed near total downregulation of DDX3X expression (Fig. 1B). On the other hand, all PCR-positive clones of Daoy cells did not show a decrease in the levels of DDX3X expression (Fig. 1B). Upon sequencing, DDX3X knockout clones of HEK293FT cells showed frameshift indels at the PAM site (Fig. 1C). However, Daoy clones showed indels in multiples of 3 nucleotides that did not result in a frameshift alteration (Fig. 1C). Thus, although the CRISPR-cas9 could successfully introduce insertion or deletion at the targeted PAM sites, only those clones carrying non-frameshift indels were viable in the case of Daoy cells. Thus, DDX3X expression appears to be essential for the viability of Daoy medulloblastoma cells.

**Figure 1.**
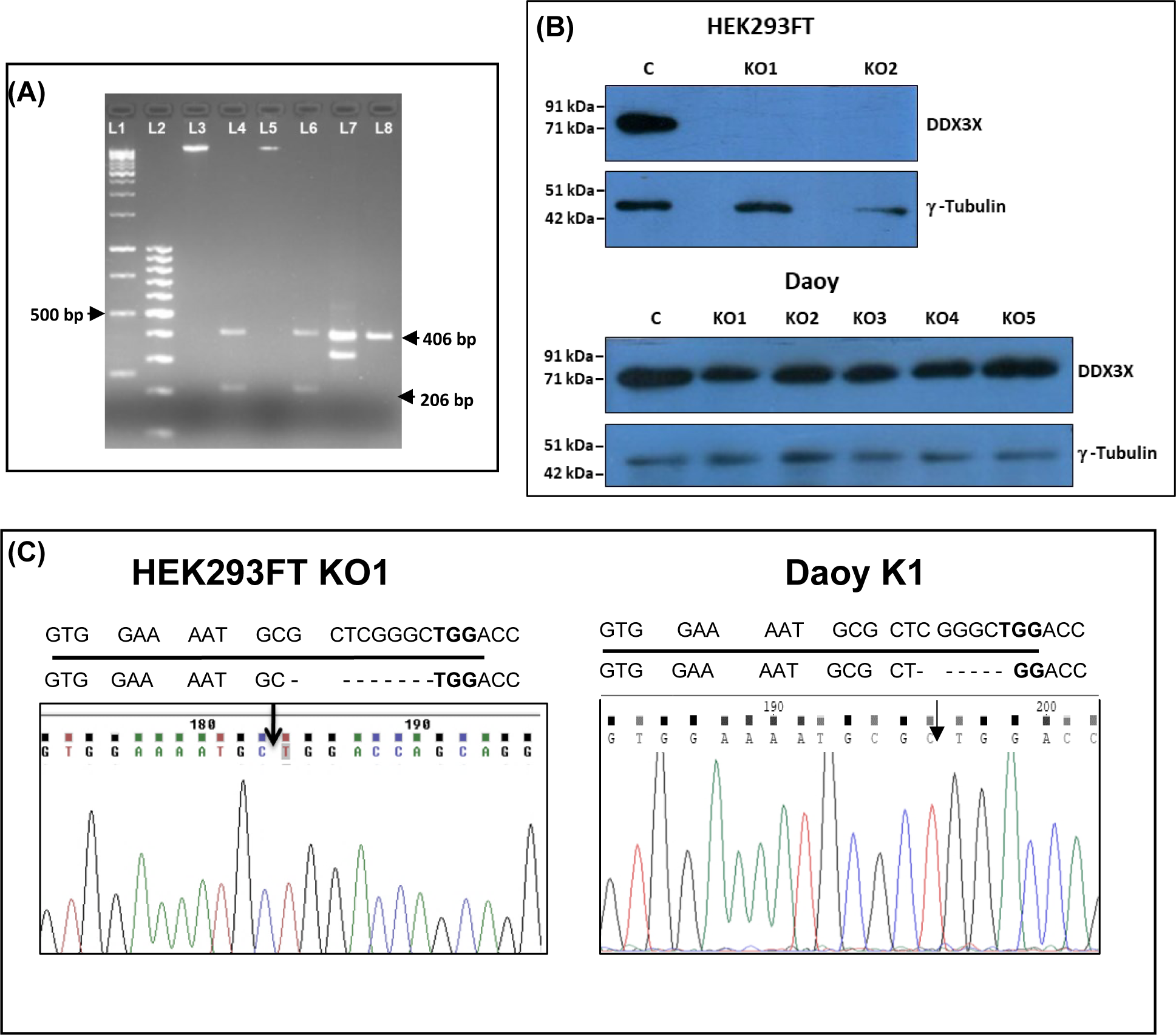
CRISPR-Cas9 mediated DDX3X knockout in HEK293FT and Daoy cells. (A) depicts the PCR-based strategy of identifying indels at the targeted PAM site. The presence of 406 bp amplicon shows success of the PCR reaction while the absence of 206 bp amplicon indicates indel at the PAM site. L1, L2: molecular size marker lanes; L3-L8: PCR products (B) Western blot analysis of the DDX3X expression in the knockout clones identified by the PCR-based strategy. γ-tubulin was used as a housekeeping loading control. (C) Representative electropherograms of Sanger sequencing of the PCR products showing the presence of 8 bp indel and 6 bp indel in the targeted PAM site in the HEK293FT and Daoy cells, respectively. The sequence underlined belongs to the guide RNA used and TGG in bold letters is the targeted PAM site.

### Knockdown of DDX3X expression inhibits proliferation and clonogenic potential of medulloblastoma cells

Since DDX3X expression could not be knocked out in the Daoy medulloblastoma cells, it was downregulated using two independent shRNAs. Oligonucleotides corresponding to shRNA sequences, one targeting the open reading frame (ORF) and the other targeting the 3’-untranslated (UTR) region were cloned in a doxycycline-inducible lentiviral vector Tet-pLKO-puro. Daoy medulloblastoma cells were transduced with the lentiviral particles carrying the shRNA construct and stably transduced cells were selected in the presence of puromycin. Western blot analysis showed a substantial reduction in the expression of DDX3X by the UTR and ORF targeting shRNA (Fig. 2A).

**Figure 2.**
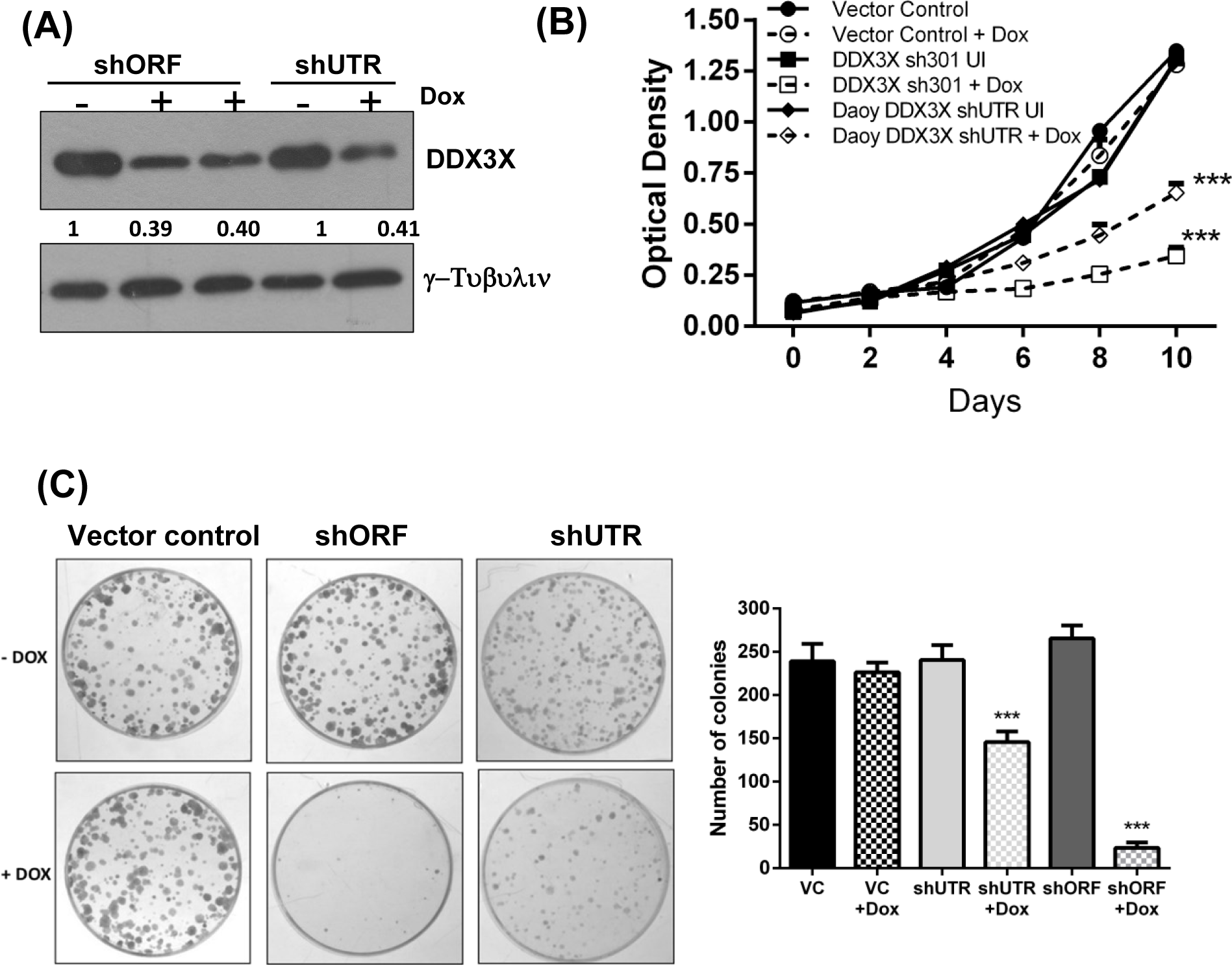
Downregulation of DDX3X expression using shRNA and its effect on the growth of the medulloblastoma cells. (A) Western blot analysis of the DDX3X expression in the two DDX3X knockdown polyclonal populations of Daoy cells before and after induction with doxycycline, using two independent shRNAs, one targeting the ORF (shORF)and the other targeting 3’-UTR (shUTR) of DDX3X. γ-tubulin was used as a housekeeping loading control. (B). Growth curves of the Daoy vector control cells and the two DDX3X knockdown populations before and after induction with doxycycline. (C) Depicts images of the plates of the clonogenic assay and the graph depicts the number of colonies formed in the clonogenic assay of the vector control (VC) and the two DDX3X knockdown populations of Daoy cells with or without doxycycline (+dox) induction *** denotes p < 0.001

The effect of DDX3X knockdown on the proliferation of the medulloblastoma cells was studied by the MTT assay. There was about a 50% to 75% decrease in the growth of Daoy cells upon the doxycycline-induced expression of the DDX3X shRNAs (Fig. 2B). There was a 50% to 95% decrease in the clonogenic potential of Daoy cells upon DDX3X knockdown compared to the vector control cells as evaluated by the clonogenic assay (Fig. 2C). The expression of the shRNA targeting the open reading frame of DDX3X, resulted in almost complete loss of clonogenic potential of the cells. Thus, DDX3X appears to be an essential gene for the survival of Daoy medulloblastoma cells as the CRISPR-cas9 mediated DDX3X knockout also failed to generate viable clones.

### Expression of DDX3X mutants restores the clonogenic and anchorage-independent growth potential of DDX3X knockdown medulloblastoma cells

The majority of mutations in the *DDX3X* gene in medulloblastomas are single nucleotide missense mutations that do not disrupt the reading frame. DDX3X possesses an ATP-dependent RNA helicase activity. Some of these missense mutations have been reported to disrupt RNA-stimulated ATP hydrolysis (15). DDX3X mutants G302V and G325E are defective in the RNA-stimulated ATPase activity. On the other hand, some of the missense mutants have been reported to potentiate TOP-FLASH reporter activity in the presence of mutant beta-catenin suggesting potentiation of the canonical WNT signaling (12). Two such mutants R534H and P568L were, therefore, selected for evaluating the effect of these mutants on medulloblastoma cell growth in the DDX3X knockdown background. Mutations were introduced in the full-length DDX3X cDNA by the site-directed mutagenesis. The wild-type and mutant DDX3X cDNAs were cloned in the p3XFLAG-CMV™-10 vector and transfected into the Daoy cells stably expressing 3’-UTR targeting DDX3X shRNA. Figure 3A shows the expression of FLAG-tagged wild-type and mutant DDX3X in the DDX3X knockdown Daoy medulloblastoma cells. The expression of wild-type or mutant DDX3X significantly increased the number of colonies formed in the clonogenic assay, almost completely rescuing the decrease in the clonogenic potential upon DDX3X knockdown (Fig. 3B, 3C). The anchorage-independent growth potential of the UTR targeting shRNA-mediated DDX3X knockdown cells decreased by about 50%, which was also restored upon the expression of wild-type as well as all the three DDX3X mutants as evaluated by the soft agar colony forming assay (Fig.3D).

**Figure 3.**
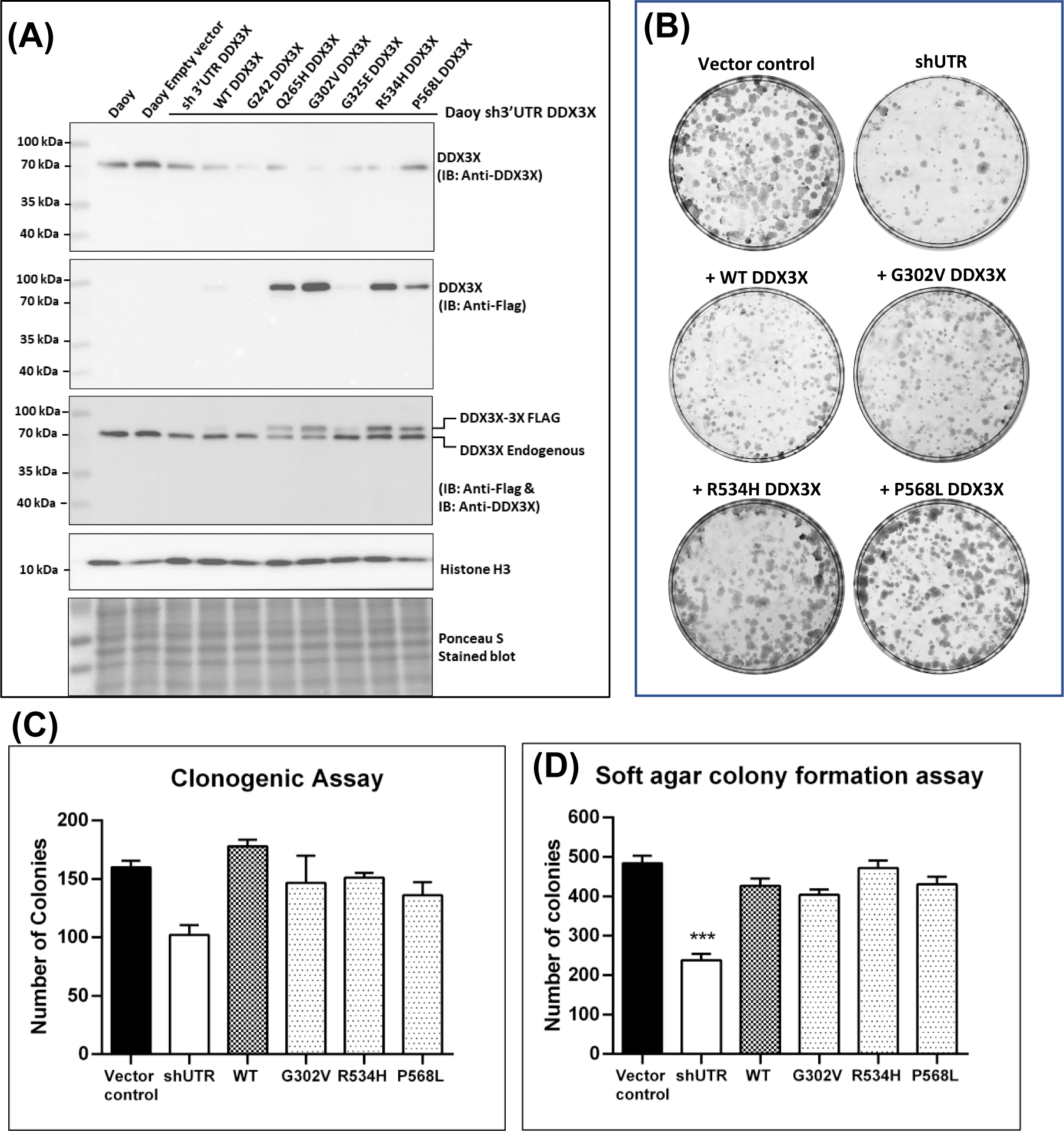
Exogenous expression of wild-type and mutant DDX3X in the shUTR mediated DDX3X knockdown population of Daoy cells and its effect on clonogenic potential and anchorage-independent growth. (A) western blot analysis of DDX3X expression in the parental, vector control, shUTR mediated knockdown population (shUTR) and the stably expressed FLAG-tagged wild-type and the indicated mutant DDX3X in the shUTR population of Daoy cells. The blot was probed with an anti-DDX3X antibody for the expression of endogenous DDX3X and an anti-FLAG antibody for the exogenously expressed DDX3X. Histone H3 was used as a housekeeping loading control. (B) shows representative images of the colonies formed in the clonogenic assay and (C), (D) depict the graph of the number of colonies formed by the vector control, shUTR, and the populations expressing exogenous DDX3X in clonogenic assay and soft agar colony forming assay, respectively. *** indicates p < 0.001

### Missense mutations in DDX3X disrupt the RNA helicase activity-dependent translation of Cyclin E having complex 5’-UTR

Exome sequencing of WNT subgroup medulloblastomas identified a rare frame-shift mutation G242fs that results in a truncated protein of 242 amino acids lacking both the RecA-like domains required for the RNA helicase activity of DDX3X (13). To check if the common missense mutations in medulloblastoma also disrupt the helicase activity, the effect of these mutations on the RNA helicase activity was investigated by studying the ability of the DDX3X mutants to unwind complex 5’-UTR of cyclin E in a luciferase reporter assay. Wild-type and mutant DDX3X cDNA were transiently transfected in a DDX3X knockout clone of the HKE293FT cells along with luciferase reporter construct carrying cyclin E 5’-UTR at the N-terminal end of the luciferase cDNA. Figure 4A shows the western blot analysis of DDX3X expression in the HEK293FT DDX3X knock-out clone before and after transfection with wild-type or mutant DDX3X cDNA. The cyclin E-5’UTR-luciferase reporter activity increased upon the exogenous expression of wild-type DDX3X but did not increase or even decreased upon the expression of the DDX3X mutants (Fig. 4B). Thus, the G242fs mutant that lacks helicase domain, G302V, G325E which are known to be defective in RNA-dependent ATP hydrolysis as well the mutants Q265H, R534H and P568L failed to promote translation from the complex 5’-UTR of cyclin E. Thus, the medulloblastoma associated missense mutants of DDX3X are defective in promoting translation from complex UTR, most likely due to defective RNA helicase activity.

**Figure 4.**
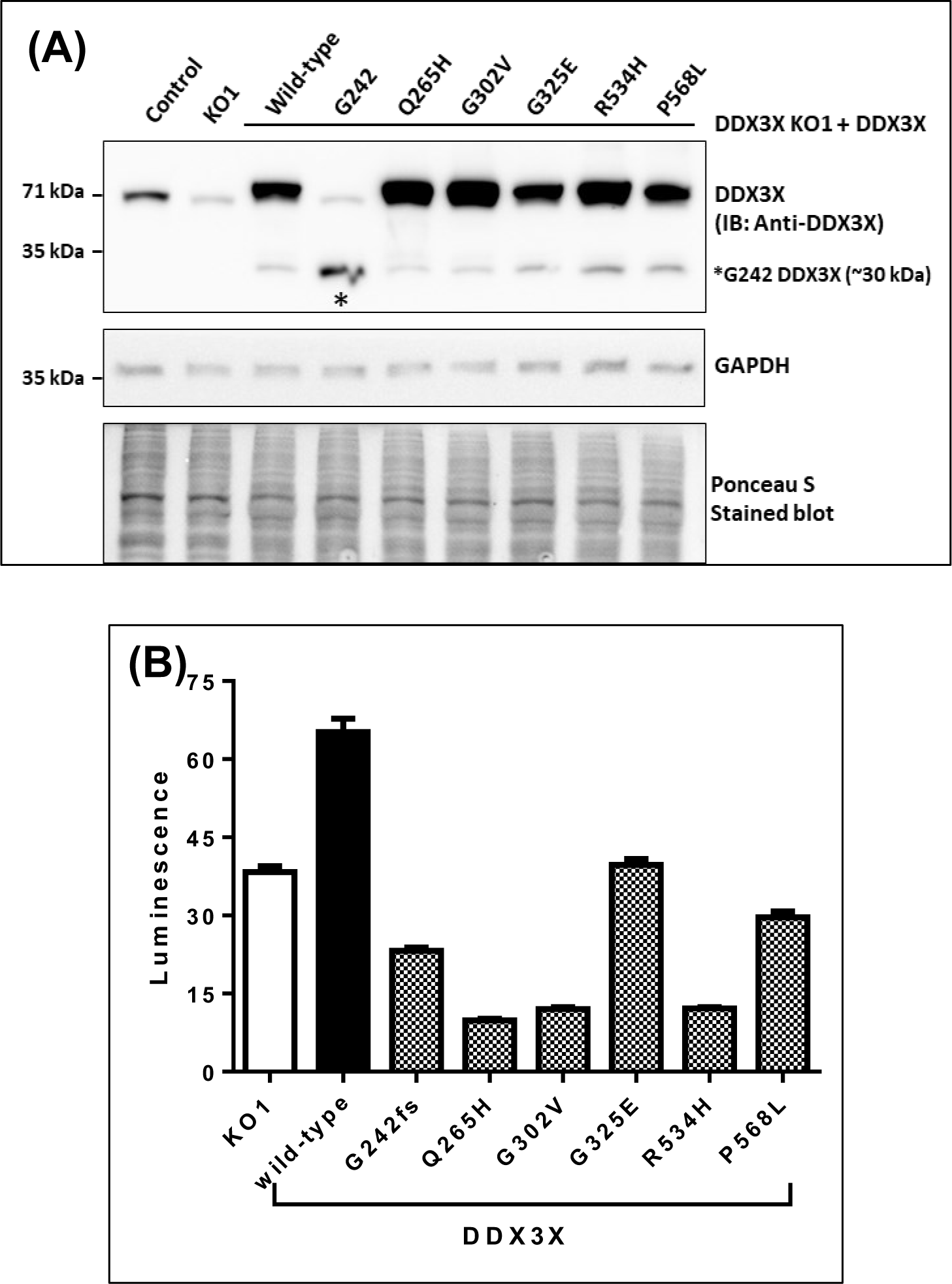
Evaluation of helicase activity-dependent translation promoting activity of DDX3X mutants evaluated by the luciferase reporter assay. DDX3X knockout clone of HEK293FT cell line was transiently transfected with the cyclin E-UTR-luciferase reporter vector, GFP expression construct, and FLAG-tagged wild-type or DDX3X mutant. (A) Western blot analysis of the expression levels of exogenously expressed wild-type or indicated mutant DDX3X probed with anti-FLAG antibody and anti-DDX3X antibody. (B) Luminescence of the luciferase reporter in the HEK293FT knockout clone cells before and after exogenous expression wild-type or mutant DDX3X. The luminescence was normalized with the GFP fluorescence to account for variation in the transfection efficiency.

### Both wild-type and mutant DDX3X participate in the stress granule formation

DDX3X is known to play a crucial role in stress granule formation (21). Therefore, stress granule formation by the DDX3X mutants was studied in the DDX3X knockdown Daoy cells. For this purpose, the cells were treated with Arsenic trioxide to induce stress granule formation. Stress granules and the exogenous wild-type and mutant DDX3X were detected using anti-eIF4G antibody and anti-FLAG antibody, respectively. Figure 5A shows stress granules formed in the Daoy cells. The FLAG-tagged wild-type as well as the three DDX3X mutants G302V, R534H, P568L localized to the stress granules formed as seen by the colocalization of anti-FLAG and anti-eIF4G fluorescent staining. Thus, the DDX3X mutants participate in stress granule formation.

**Figure 5.**
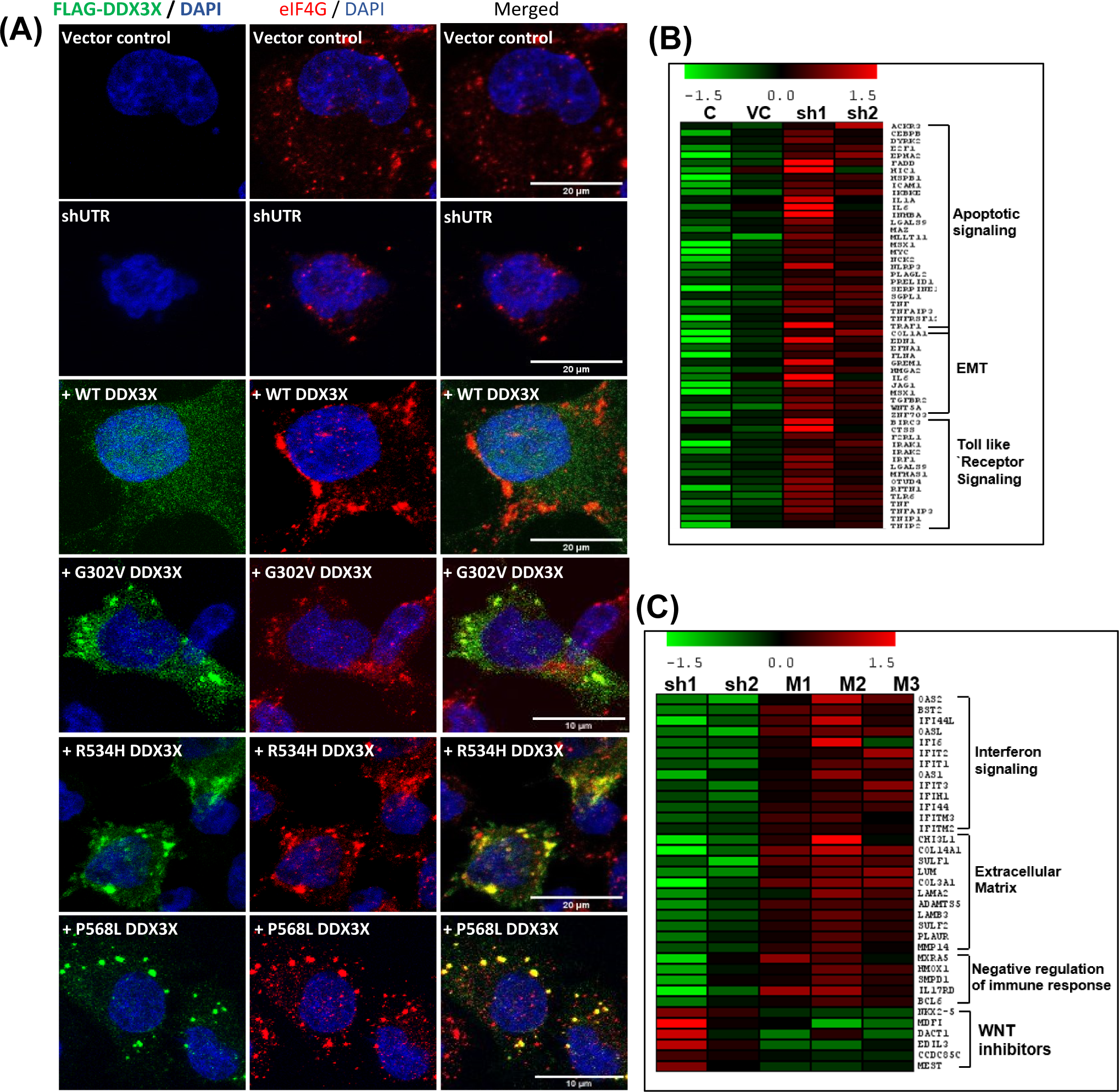
Participation of exogenously expressed wild-type and mutant DDX3X in the stress granule formation in the DDX3X knockdown medulloblastoma cells and Heatmaps of genes significantly differentially expressed upon DDX3X knockdown and on the expression of wild-type and mutant DDX3X in DDX3X knockdown Daoy cells. (A) Daoy cells expressing DDX3X shRNA (shUTR) and those shUTR cells stably expressing wild-type and mutant DDX3X were treated with Arsenic trioxide for induction of stress granules. The stress granules were detected by confocal imaging of immunofluorescence using anti-eIF4G antibody while the exogenously expressed FLAG-tagged DDX3X was detected using anti-FLAG antibody. Heat-maps of the genes significantly upregulated (B) upon the shRNA-mediated DDX3X knockdown medulloblastoma cells and the genes significantly upregulated (C) upon the expression of mutant DDX3X in the DDX3X knockdown cells compared to the knockdown cells in the indicated pathways. C:parental Daoy cells; VC:vector control; sh1, sh2:shRNA expressing cells; M1,M2,M3: DDX3X mutants G320V, R534H and P568L.

### Transcriptome sequencing shows downregulation of interferon signaling, upregulation of immune system activation genes upon DDX3X knockdown, and restoration of their expression by the mutant DDX3X

Transcriptome sequencing of the parental Daoy cells, vector control population, as well as both DDX3X ORF and UTR targeting shRNA expressing polyclonal populations, was carried out using the Illumina HiseqX platform. Several pathways like activation of the stress-activated MAPK pathway, immune system development and activation, Toll-like receptor signaling, inflammatory response, and apoptotic signaling were enriched (p_adj_ < 0.05) in the genes significantly upregulated upon DDX3X knockdown in the medulloblastoma cells (Fig. 5B, 6A). Upregulation of these pathways could be responsible for the loss of viability upon complete loss of DDX3X expression. On the other hand, pathways like epithelial-mesenchymal transition, cell migration, cell adhesion which were also upregulated could increase malignant potential upon DDX3X knockdown in the medulloblastoma cells (Fig. 6A). The interferon signaling, insulin-like growth factor signaling, regulation of cell growth pathways were enriched (p_adj_ < 0.05) in the genes significantly downregulated consistent with the DDX3X knockdown leading to the growth inhibition of medulloblastoma cells (Fig. 6B).

**Figure 6.**
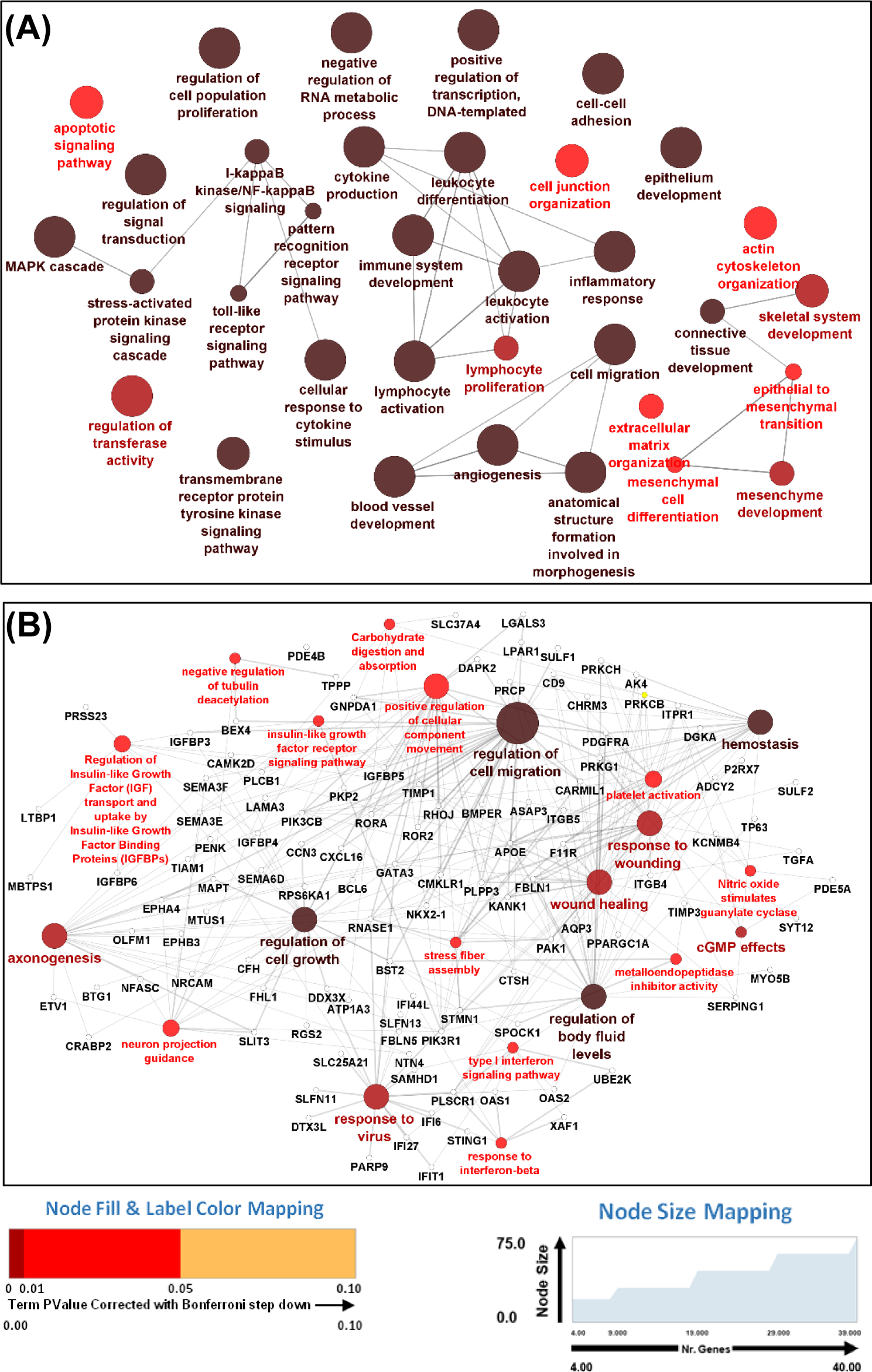
Biological pathways and protein-protein interaction network of the genes significantly differentially expressed upon DDX3X knockdown. (A) Biological pathways enriched significantly upregulated upon DDX3X knockdown in Daoy cells (B) Biological pathways enriched and protein-protein interaction network of the genes significantly downregulated upon DDX3X knockdown in Daoy cells. padj = Bonferroni correction adjusted p-value.

Transcriptome sequencing of Daoy cells expressing DDX3X 3’-UTR targeting shRNA and stably transfected with wild-type or mutant DDX3X was also done. Pathways like interferon signaling pathway, extracellular matrix proteins, cholesterol metabolic process, and negative regulation of molecular mediators of immune response were enriched among the genes significantly upregulated upon the expression of mutant DDX3X (Fig. 5C, 7). Thus, the expression of interferon signaling pathway genes which decreased upon the DDX3X knockdown was restored upon the expression of mutant DDX3X. Upregulation of negative regulators of immune system activation could overcome the activation of immune response-mediated loss of viability upon DDX3X knockdown. Besides, upregulation of the genes belonging to the pathways like extracellular matrix proteins, cholesterol metabolic process, and decrease in the levels of negative regulators of the WNT signaling pathway (Fig. 5C, Fig. 7) could increase the malignant potential of the medulloblastoma cells upon the expression of mutant DDX3X.

**Figure 7.**
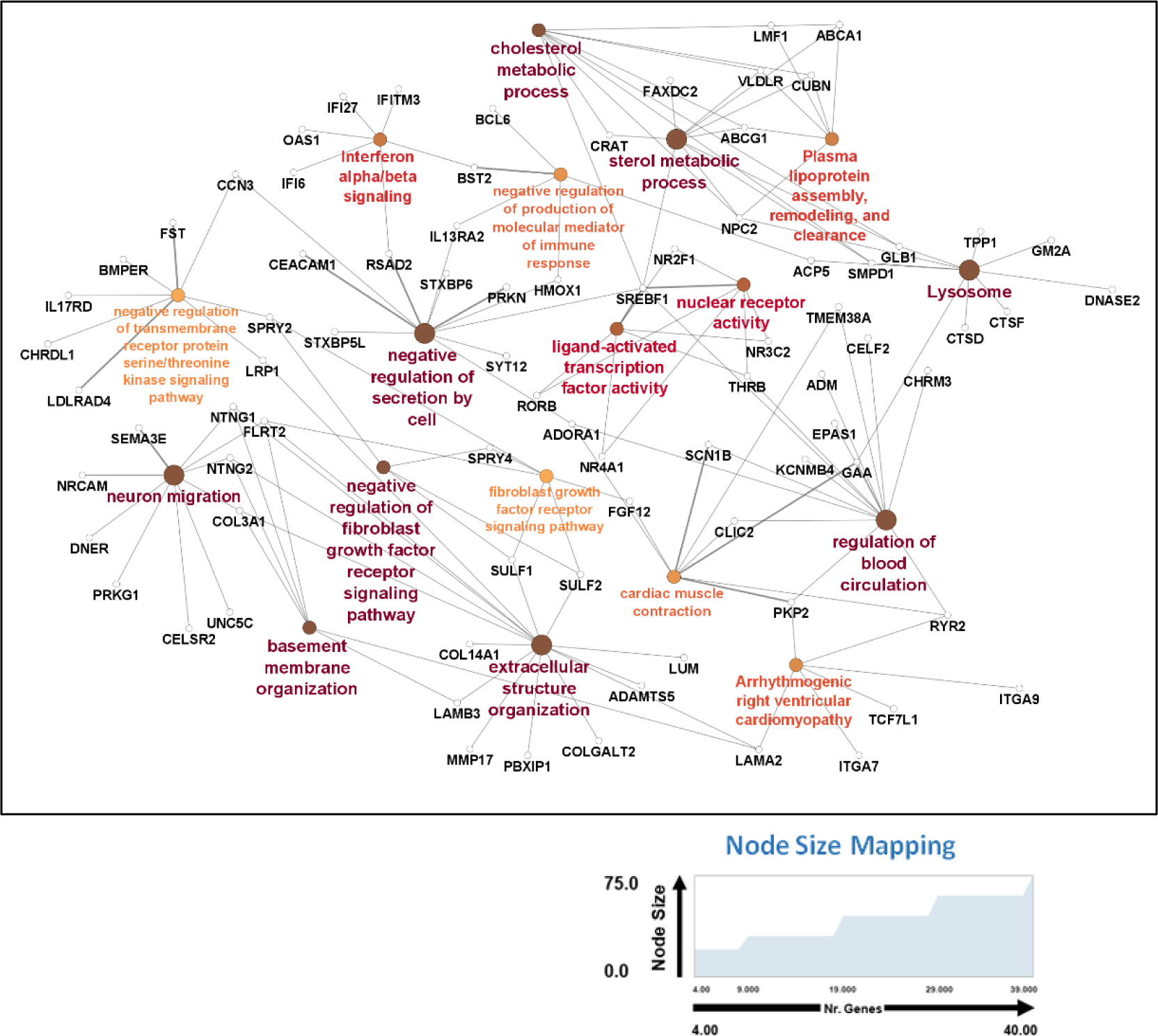
Biological pathways significantly enriched in the genes upregulated upon expression of DDX3X mutants in the DDX3X knockdown medulloblastoma cells. The figure shows the interaction network of the pathways significantly enriched (p_adj_ < 0.05) in the genes upregulated upon the expression of the DDX3X mutants compared to the shUTR-mediated DDX3X knockdown Daoy cells. padj = Bonferroni correction adjusted p-value.

## Discussion

### DDX3X expression is essential for the viability, growth, and malignant potential of the medulloblastoma cells

DDX3X being a multifunctional RNA helicase is known to play a complex role in the pathogenesis of several cancers (6). Mutations in DDX3X are restricted to medulloblastoma subgroups driven by the canonical WNT and SHH signaling pathways (10). The present study was carried out using Daoy, a SHH subgroup cell line. CRISPR-cas9 mediated DDX3X knockout clones could not be generated in the medulloblastoma cells while the same strategy was successful in generating DDX3X knockout clones of HEK293FT cells. The shRNA-mediated DDX3X knockdown also resulted in considerable growth inhibition and almost complete loss of clonogenic and anchorage-independent growth potential of the medulloblastoma cells. These findings are consistent with the fact that almost all mutations reported in medulloblastomas are non-truncating mutations suggesting DDX3X expression is essential for the viability of medulloblastoma cells.

### Helicase activity of DDX3X is dispensable for the viability of the medulloblastoma cells

Most DDX3X are missense or non-truncating indel mutations in medulloblastoma located within the two RecA-like domains. These domains contain the ATPase and RNA helicase motifs required for the ATP hydrolysis and RNA substrate binding and unwinding (11,12,14). Several DDX3X mutants in medulloblastoma are reported to be defective in the ATPase activity that is essential for its ATP-dependent helicase activity (15). The helicase activity of DDX3X is known to be required for its role in translation particularly of mRNAs having complex 5’-UTRs like cyclin E (22,23). All the medulloblastoma-associated DDX3X mutants tested were found to be defective in the unwinding of the complex 5’-UTR of cyclin E. Three medulloblastoma-associated DDX3X mutants R276K, R276Y, and R543H which could be tested in vitro using DDX3X 132-607 construct were found to be defective in helicase activity (16). Thus, the medulloblastoma-associated DDX3X mutants appear to be defective in their helicase activity. Exogenous expression of these DDX3X mutants could restore the clonogenic potential and anchorage-independent growth potential of the DDX3X knockdown medulloblastoma cells. Thus, while the complete loss of expression of DDX3X is lethal, the helicase defective mutants could restore the viability and malignant potential of the medulloblastoma cells. Therefore, the helicase activity of DDX3X appears to be dispensable for the viability of the medulloblastoma cells.

### The N-terminal domain of medulloblastoma-associated DDX3X mutants is functional, participates in stress granule formation, and could thereby restore the viability of the medulloblastoma cells lost upon loss of DDX3X expression

Transcriptome analysis of DDX3X knockdown medulloblastoma cells showed upregulation of genes involved in the stress-activated MAPK signaling, apoptotic signaling pathway, and immune system activation including the Toll-like receptor signaling pathway suggesting cells are under stress upon the loss of DDX3X expression. Under stress, global translation stops, and non-translating mRNAs and proteins assemble to form stress granules that allow cells to survive. DDX3X is known to play a critical role in stress granule formation. Downregulation of DDX3X has been shown to interfere with stress granule assembly, thereby reducing cell viability upon stress (21). Medulloblastoma-associated DDX3X mutants were found to participate in the stress granule formation and hence could overcome the loss of viability and clonogenic potential brought about by the loss of DDX3X expression.

The stress granule-inducing capacity of DDX3X is known to be independent of its ATPase, helicase activity and maps to its N-terminal region. Similarly, DDX3X’s role in the interferon signaling maps to its N-terminal domain. DDX3X facilitates phosphorylation of the interferon regulatory factor 3 by IKKα and IKKε for which phosphorylation of amino acid residues in its N-terminal domain is required (23,24). The medulloblastoma-associated DDX3X mutants could also restore the expression of the genes involved in the interferon signaling pathway as they have the N-terminal domain intact. Thus, the medulloblastoma-associated DDX3X mutants are defective in their helicase activity but have their N-terminal domain intact and functional which overcomes the lethality induced by the loss of DDX3X expression.

### Translation promoting activity of DDX3X appears to play the role of a tumor suppressor in the pathogenesis of medulloblastoma

In the present study, all the DDX3X mutants tested were found to be defective in the helicase activity-dependent translation-promoting activity of DDX3X. These DDX3X mutants upregulated the expression of several genes involved in the malignant potential like extracellular matrix and cholesterol metabolism consistent with the upregulation of the clonogenic and anchorage-independent growth potential upon their expression in the medulloblastoma cells. Thus, DDX3X appears to act as a tumor suppressor in the medulloblastoma cells. In DDX3X heterozygous and homozygous conditional knockout conditions, there was a significant increase in the tumor incidence and severity in the WNT and SHH medulloblastoma mouse models (25). Thus, DDX3X acted as a tumor suppressor in these mouse models as well.

How the helicase activity of DDX3X acts as a tumor suppressor in medulloblastoma cells is not understood. DDX3X is known to activate casein kinase and thereby stimulate canonical WNT signaling pathway (26,27) and this kinase activation is reported to be mutually exclusive with its ATPase activity. Hence, the ATPase/helicase defective DDX3X mutants have been suggested to act as constitutively active mutants having the kinase activating function. The casein kinase activating function maps to the C-terminal domain of DDX3X (27). Therefore, rare truncating mutations in DDX3 will not be able to activate the WNT signaling by this mechanism. In the WNT subgroup medulloblastomas, wherein activating mutations in the beta-catenin encoding gene stabilize the beta-catenin expression levels, the decrease in the levels of negative nuclear regulators of the canonical WNT signaling could be the primary role of mutant DDX3X in the stimulation of WNT signaling.

How the loss of helicase activity upregulates the malignancy-associated genes like extracellular matrix genes, cholesterol synthesis genes and downregulates the negative regulators of the WNT signaling pathways remains to be determined. Cancer-associated DDX3X mutants have been reported to impair global translation and drive stress granule hyper-assembly in Hela cells (28). However, in this study DDX3X mutants were overexpressed in the presence of endogenous DDX3X, and overexpression of wild-type DDX3X also led to stress granule assembly. On the other hand, medulloblastoma-associated DDX3X mutants were found to cause specific defects in translation in Ded1-mutant yeast cells (29). The DDX3X mutants were found to be defective in the translation of mRNAs having secondary structures in their 5’-UTR. Identification of the mRNAs with complex 5’-UTR whose translation is affected by the expression of DDX3X mutants would reveal how the loss of helicase activity increases the expression of malignancy-associated genes and decreases the expression of negative regulators of WNT signaling.

### Dual role of DDX3X in the pathogenesis of medulloblastoma

Most medulloblastoma-associated DDX3X mutants are non-truncating missense or indels which appears to be inconsistent with its role as tumor suppressor. DDX3X expression was found to be essential for the viability of the medulloblastoma cells consistent with the non-truncating nature of the DDX3X mutations. However, the helicase-defective DDX3X mutants could rescue the lethality and loss of malignant potential. The N-terminal domain having a crucial role in stress granule formation, interferon signaling was found to be intact and functional in these mutants. Therefore, although the helicase activity of DDX3X appears to act as a tumor suppressor, its N-terminal domain is essential for the viability of the medulloblastoma cells for being a crucial mediator of the stress granule formation. Thus, N-terminal domain of DDX3X appears to act as an oncogene while the translation promoting activity as a tumor suppressor in the pathogenesis of medulloblastoma.

### DDX3X as a therapeutic target in medulloblastoma

Recurrent mutations in U1 spliceosomal small nuclear RNAs are known to occur in about 50% of adult SHH medulloblastomas which could elevate the expression of cryptic transcripts (30). Proteome analysis has found higher expression of RNA surveillance pathways in adult SHH medulloblastomas (31). Interestingly, DDX3X mutations are common, particularly in adult SHH subgroup medulloblastomas. DDX3X knockdown has been reported to result in the accumulation of cellular double-stranded RNAs triggering innate immune response mediated cell death in breast cancer cells (32). Therefore, DDX3X could serve as a therapeutic target in adult SHH medulloblastomas. Inhibitors of the N-terminal domain of DDX3X and not inhibitors of its helicase activity could have therapeutic potential in the treatment of WNT and SHH subgroup medulloblastomas.

## Acknowledgments

We would like to acknowledge technical assistance from Sadaf Shaikh, Umesh Kadam, and Anant Sawant

