## Supplementary Table 1 for "Medulloblastoma-associated DDX3X mutants are oncogenic having a defect in translation-promoting activity but functional in stress granule formation and interferon signaling"

**List of oligonucleotide sequences used**

| **Name** | **Forward (5’-3’)** | **Reverse (5’-3’)** |
| --- | --- | --- |
| DDX3X_CRISPR_gRNA1 | CACCGAGTGGAAAATGCGCTCGGGC | AAACGCCCGAGCGCATTTTCCACTC |
| DDX3X_CRISPR_gRNA2 | CACCGGATCTCGTAGTGATTCAAGA | AAACTCTTGAATCACTACGAGATCC |
| DDX3X_KO_PCR Screen | F1- GAATTGCGGTGTGAGAGGGA  F2- AGTGGAAAATGCGCTCGGG | GACAGCCCCAACCAACATTC |
| DDX3X_shORF | CCGGCGGAGTGATTACGATGGCATTCTCGAGAATGCCATCGTAATCACTCCGTTTTT | AATTAAAAACGGAGTGATTACGATGGCATTCTCGAGAATGCCATCGTAATCACTCCG |
| DDX3X_shUTR | CCGGCCCTGCCAAACAAGCTAATATCTCGAGATATTAGCTTGTTTGGCAGGGTTTTT | AATTAAAAACCCTGCCAAACAAGCTAATATCTCGAGATATTAGCTTGTTTGGCAGGG |
| DDX3X_cDNA_WT | GGTGGATCCATGAGTCATGTGGCAGTGGAA | GTCGAATTCCTATCAGTTACCCCACCAGTCAAC |
| DDX3X_SDM_G242fs | GAAGGAAAATGGAGGTATGGGCGCCG | CGGCGCCCATACCTCCATTTTCCTTC |
| DDX3X_SDM_Q265H | GAGTTGGCAGTACACATCTACGAGGAAGC | GCTTCCTCGTAGATGTGTACTGCCAACTC |
| DDX3X_SDM_R534H | GTATTGGTCGTACGGGACATGTAGGAAACCTTGGC | GCCAAGGTTTCCTACATGTCCCGTACGACCAATAC |
| DDX3X_SDM_P568L | GAAGCTAAACAAGAAGTGCTGTCTTGGTTAGAAAACATGG | CCATGTTTTCTAACCAAGACAGCACTTCTTGTTTAGCTCT |
| CCNE1_5UTR | GATAAGCTTTCTGAGCCGGGCGCAGGA | R1- ATGGGGCTGCTCCGGCC  R Nested- GACAAGCTTGATGGGGCTGCTCCGGCCTGAGGCCAGGGTCTTGTCCGCGGC |
